# Design to Data for mutants of β-glucosidase B from *Paenibacillus polymyxa*: N160L, N160S, N160C, N160M, N160G

**DOI:** 10.1101/2025.11.24.690255

**Authors:** Nicholas Li, Ashley Vater, Justin B. Siegel

## Abstract

Protein design is advancing toward quantitative modeling of enzyme function and stability. However, progress remains limited by the scarcity of standardized experimental datasets for training and benchmarking computational models. The Design to Data (D2D) program addresses this need by generating harmonized measurements of catalytic and stability parameters across an extensive β-glucosidase B (BglB) variant library. Here, we expand the D2D dataset with kinetic and thermal characterization of five single-point BglB variants and the wild-type (WT), including soluble expression, Michaelis-Menten constants (*k*_cat_, K_M_, and *k*_cat_/K_M_), and melting temperature (T_M,_). Foldit Standalone was used to model the structural effects of the mutations. In this study, a weak but consistent association between Foldit total system energy (TSE) and T_M_ was observed, suggesting local energetic effects that may influence stability. Together with the broader D2D corpus, these data enhance the functional mapping of BglB and provide model-ready benchmarks for developing and evaluating data-driven predictors of enzyme activity and stability.

## INTRODUCTION

The advances in artificial intelligence are catalyzing a revolution in the biotechnology field ^1^. In this domain, rational enzyme modification enables catalytic functions not found in nature ^2–5^, providing a foundation for developing next-generation protein therapeutics, biofuels, food science, and bioremediation ^6–8^. The intersection of artificial intelligence and enzyme research underscores the growing need for protein design software that predicts function, building off the developments around structure prediction capabilities. However, continued breakthroughs in this field are limited by the availability of large, structured, and interoperable datasets ^9^.

These dataset limitations are particularly relevant to the enzyme space, given the nuanced and complicated nature of the reactions they catalyze. Specifically, the field lacks comprehensive datasets containing quantitative measures of catalytic activity (*k*_cat_, K_M_, *k*_*cat*_/K_M_) and protein unfolding temperature (functional thermal stability, T_M_) ^10^. To address this challenge, national undergraduate research network Design to Data (D2D) was established in 2019 to systematize dataset expansion. This program enables students to measure and record quantitative kinetic and thermal stability data for single-point mutations of β-glucosidase B (BglB) ^11^. Program coordination is based at the University of California, Davis, and has led to the dataset’s growth through contributions from a national consortium of more than 40 institutions. Previous findings by Carlin et al. revealed the current predictive capabilities of various computational methods tested on a dataset of 100 kinetically characterized BglB single-point mutants, demonstrating weak relationships between experimental data and structural metrics such as interface energy ^10^. Further analysis showed that T_M_ and algorithm-derived features (Rosetta and FoldX) also exhibited weak correlations ^12^. Collectively, the authors of these studies recommend that expanding the BglB dataset may have utility in improving predictive accuracy of newer models.

This study analyzed the kinetic properties and thermal stability of five novel single-point mutations (N160L, N160S, N160C, N160M, and N160G) in BglB from *Paenibacillus polymyxa*. Although residue N160 does not directly contact the ligand, its proximity to the active site suggests potential influence through indirect mechanisms. We investigated the relationship between Foldit Standalone algorithm-derived features and both catalytic performance and thermal stability.

## METHODS

### BglB mutant design

Five BglB mutants were designed using FoldIt Standalone, an interactive graphical user interface that allows for real-time direct manipulation of protein structures ^13^. FoldIt Standalone quantifies protein structure stability using the Rosetta energy function ^2^. Mutants were assigned unique total system energy (TSE) scores and individual residue energy values subsequent to the application of both the minimization and repack/shake functions to the residue of interest and 29 neighboring residues.

### Mutagenesis

Using established protocols, the Kunkel method was used for oligonucleotide-directed mutagenesis to produce the five single-point mutants of the already cloned pET29b+ vectors ^14^. Subsequently, transformed *E. coli* DH5α colonies were selected, and plasmid DNA was extracted and purified using standard miniprep protocols. Plasmid DNA was sent for Sanger Sequencing (Genewiz, Arzenta). Sequence data was analyzed (Benchling) to verify the site mutation and confirm that the remaining sequence was error-free.

### Protein production and purification

Transformed *E. coli* BL21 cells were expanded and the BglB variants were expressed as previously described ^12^. Isopropyl β-D-1-thiogalactopyranoside (IPTG) was used to induce expression of the target proteins. The cells were lysed and the histidine tagged BglB proteins were purified with nickel-based immobilized metal affinity chromatography. Variant and WT protein concentration was determined by A280 using a BioTek Epoch spectrophotometer. Protein purity was analyzed using sodium dodecyl sulfate polyacrylamide gel electrophoresis (SDS-PAGE).

### Michaelis-Menten kinetics and thermal stability of mutants

Kinetic activities of enzyme variants were quantified by monitoring hydrolysis of 4-nitrophenyl-β-D-glucopyranoside at 420 nm, following the established protocol ^12^ with minor modifications. To enhance resolution at lower substrate concentrations, a 3× dilution series (100 mM, 33.33 mM, 11.11 mM, 3.70 mM, 1.23 mM, 0.41 mM, 0.14 mM, and 0 mM) was used in place of the original 4× series, and absorbance was recorded at 1-minute intervals for 15 minutes. The kinetic data was fitted to the Michaelis-Menten model using a SciPy nonlinear curve fitting function. *k*_cat_ and K_M_ values were calculated for each variant and the catalytic efficiency (*k*_cat_/K_M_) was reported.

Thermal stability of enzyme variants was quantified using a fluorescence-based protein unfolding assay with the Protein Thermal Shift™ (PTS) dye kit (Applied Biosystems) on a QuantStudio 3 System. Purified enzymes were diluted 2× prior to the addition of 16× PTS dye. Fluorescence was monitored as the temperature increased from 20 °C to 90 °C, and melting temperatures (T_M_) were derived from the inflection point of the two-state Boltzmann model.

For both mutant and wild-type enzymes, sample sizes for T_M_ determination were ≤3. Because the assumption of equal variances is unreliable in small samples, Welch’s t-test was used to compute test statistics and 95% confidence intervals. Pairwise intervals were constructed at an overall significance level of α = 0.05.

## RESULTS

### Mutation Selection

Using Foldit Standalone, residue N160 was selected for mutational analysis due to its proximity to the active site. Although N160 does not directly contact the ligand, it lies approximately 12 Å away (Figure 1) and contributes to the stabilization of a local residue network through hydrogen bonding with T116 and T218 (Figure 2a, yellow dotted lines). Five substitutions (leucine, serine, cysteine, methionine, and glycine) were introduced at this position to assess how differences in side-chain properties influence catalytic performance and thermal stability. Foldit Standalone predictions indicated that all five mutations disrupted the hydrogen-bonding network at N160, thereby eliminating the stabilizing interactions observed in the wild-type enzyme. Consequently, all variants were hypothesized to exhibit reduced catalytic efficiency and thermal stability relative to wild type.

**Figure 1.**
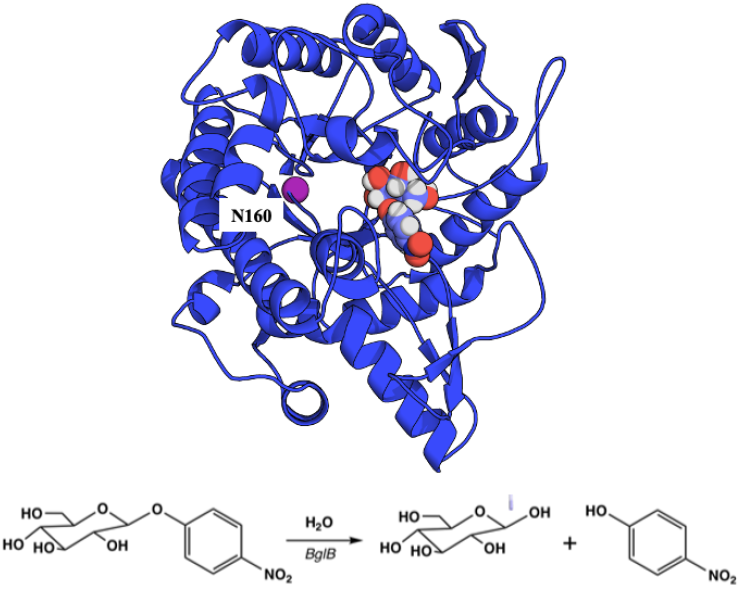
(a) Ribbon diagram of BglB (blue structure) with substrate 4-nitrophenol beta-D glucopyranoside (light indigo structure) created with PyMOL. The residue location, N160, is labeled with a magenta sphere. The ligand is labeled as multiple spheres. (b) Illustration of the BglB reaction of the reporter substrate, p-nitrophenyl-beta-D-glucoside (pPNG).

**Figure 2a.**
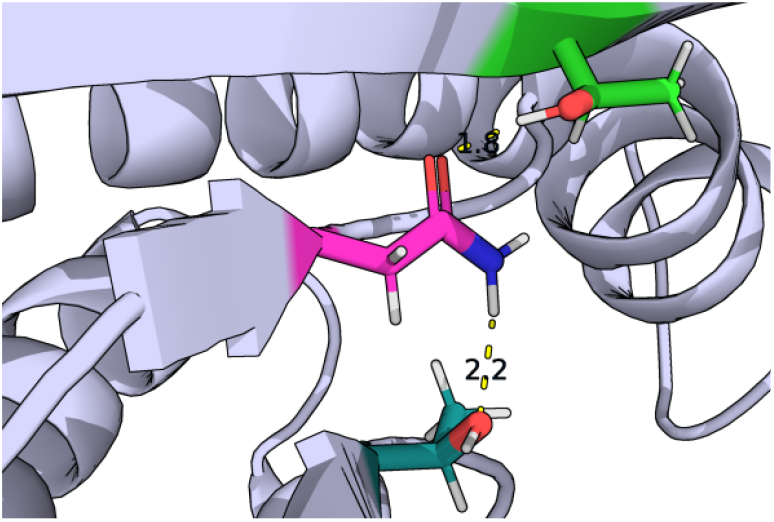
PyMOL-generated image of residue, N160 (magenta). Hydrogen bonds and distances are labeled and illustrated as yellow-dashed lines with neighboring residues, T218 (green, 1.8 Å) and T116 (dark cyan, 2.2 Å).

The leucine and methionine substitutions introduced bulkier side chains (Figure 2b), increasing the clashing score from 0.843 to 5.401 and 15.003, respectively. Both mutations also resulted in positive changes in TSE: +6.46 for leucine and +5.36 for methionine. Given the increased steric clash and elevated TSE values, these variants were predicted to exhibit reduced catalytic efficiency and thermal stability compared with the wild-type enzyme.

**Figure 2b.**
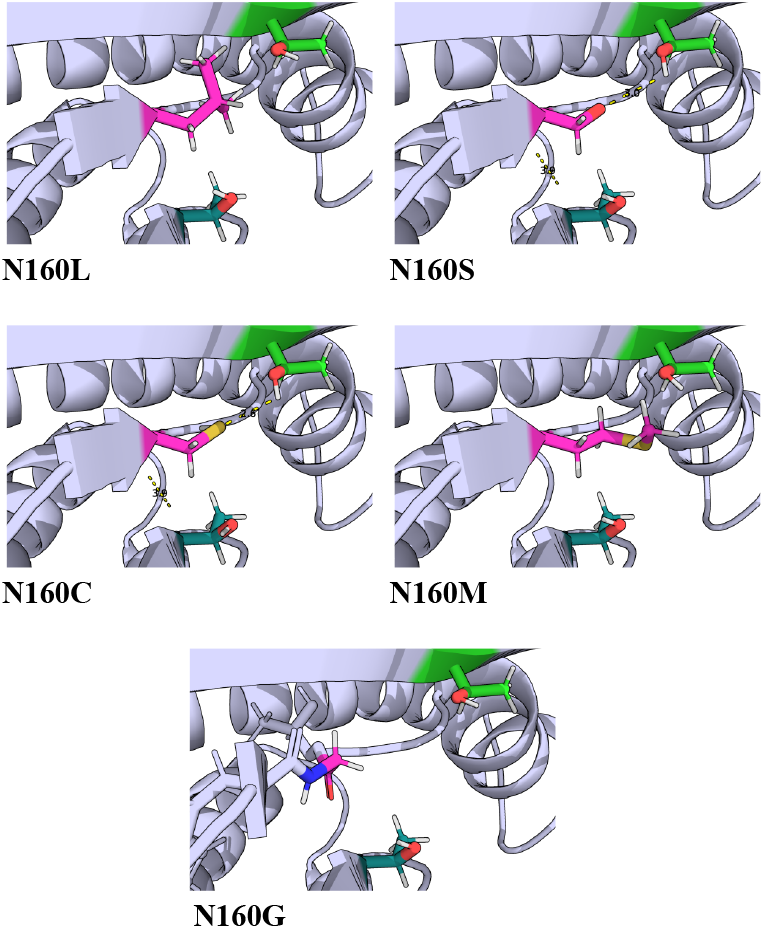
PyMOL generated images of five variant residues shown in magenta. Neighboring T218 and T116 residues are shown as light green and dark cyan, respectively. Alpha carbon distances labeled between variant residues and T218 and T116 for N160S (3.0 Å, 3.9 Å) and N160C (2.6 Å, 3.9 Å).

The mutations to cysteine and glycine introduced smaller side chains relative to asparagine (Figure 2b), with side-chain scores of 0.098 and 0, respectively, both lower than that of asparagine (2.401). Because of the reduced size and increased flexibility of these residues, the mutations were predicted to diminish catalytic efficiency, as the loss of steric constraints could impair precise substrate positioning. Both cysteine and glycine substitutions also produced positive changes in TSE of +1.64 and +11.66, respectively, suggesting decreased thermal stability. In contrast, substitution of asparagine with serine was expected to cause minimal functional change given the two residues’ similar polarity and hydrophilicity. Further the serine mutation resulted in a positive TSE shift of +1.79, indicating a negligible predicted difference in the protein’s thermodynamic folding favorability.

#### Protein Purity and Expression

All mutant proteins exceeded concentration of 1.60 mg/mL as determined by absorbance at 280 nm. SDS–PAGE analysis confirmed the presence and purity of the expressed proteins (Figure 3). Protein concentrations for all mutants and the wild-type ranged from 1.6 to 3.5 mg/mL (Table 1). The consistent and intense band patterns indicated successful expression and sufficient purity for subsequent kinetic and thermal characterization.

**Table 1.** Total system energy (TSE), protein concentration, kinetic parameters (*k*_cat_, K_M_, *k*_cat_/K_M_), and thermal stability (T_M_) of wild-type and variant BglB enzymes.

| | TSE | Expression [mg/mL] | $k_{cat}$ [$\text{min}^{-1}$] | $K_M$ [mM] | $k_{cat}/K_M$ [$\text{min}^{-1}\text{mM}^{-1}$] | $T_M$ [$^{\circ}\text{C}$] |
| --- | --- | --- | --- | --- | --- | --- |
| N160L | -1083.2 | 3.1 | $194.7 \pm 7.6$ | $13.8 \pm 1.6$ | $14.1 \pm 1.7$ | $45.2 \pm 0.3$ |
| N160S | -1087.9 | 2.8 | $271.6 \pm 3.0$ | $4.9 \pm 0.2$ | $55.2 \pm 2.4$ | $44.2 \pm 0.5$ |
| N160C | -1088.1 | 2.4 | $548.8 \pm 9.4$ | $4.3 \pm 0.3$ | $128.5 \pm 8.8$ | $46.3 \pm 0.4$ |
| N160M | -1084.3 | 1.6 | $249.7 \pm 7.3$ | $14.1 \pm 1.3$ | $16.8 \pm 1.5$ | $43.1 \pm 0.7$ |
| N160G | -1082.7 | 2.5 | $588.6 \pm 8.2$ | $3.5 \pm 0.2$ | $170.5 \pm 9.9$ | $43.0 \pm 0.5$ |
| WT | -1089.7 | 3.5 | $393.0 \pm 7.0$ | $5.0 \pm 0.3$ | $78.9 \pm 5.4$ | $45.9 \pm 0.2$ |

**Figure 3.**
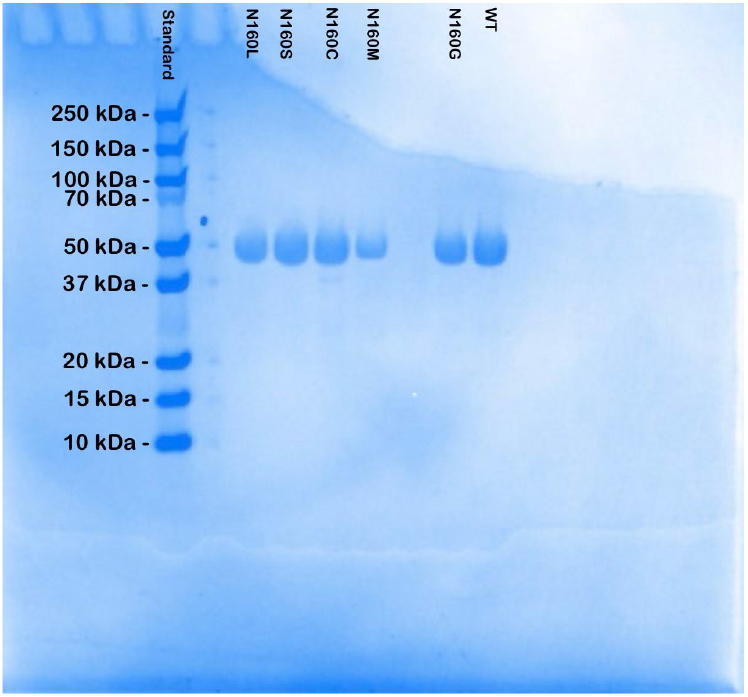
SDS-PAGE of wild-type and five variant enzymes. BglB wild-type and mutant bands formed around 50 kDA. The order of mutants and protein concentration shown from left to right: N160L, N160S, N160C, N160M, and N160G.

### Kinetic Activity

Building on the BglB dataset, we determined kinetic constants for three novel mutants: N160L, N160M, and N160G; N160S and N160C had been previously characterized but the biological replicate data presented here represents important internal control and validation measures for the larger study. Substrate saturation curves were generated for the wild-type and all variants using a standard kinetic assay (Figure 4). The wild-type enzyme exhibited a catalytic efficiency (*k*_cat_/K_M_) of 78.9 ± 5.4 min^-1^mM^-1^, consistent with previously reported values. N160C and N160G showed higher catalytic efficiencies of 128.5 ± 8.8 min^-1^mM^-1^ and 170.5 ± 9.9 min^-1^mM^-1^, respectively. In other words, N160C and N160G also had higher turnover rates (*k*_cat_) and enhanced approximate binding affinity constants (K_M_) than the wild-type. The other three mutants had reduced turnover rates (Table 1) and the K_M_ values for N160L and N160G were more than twice that of the wild-type.

**Figure 4.**
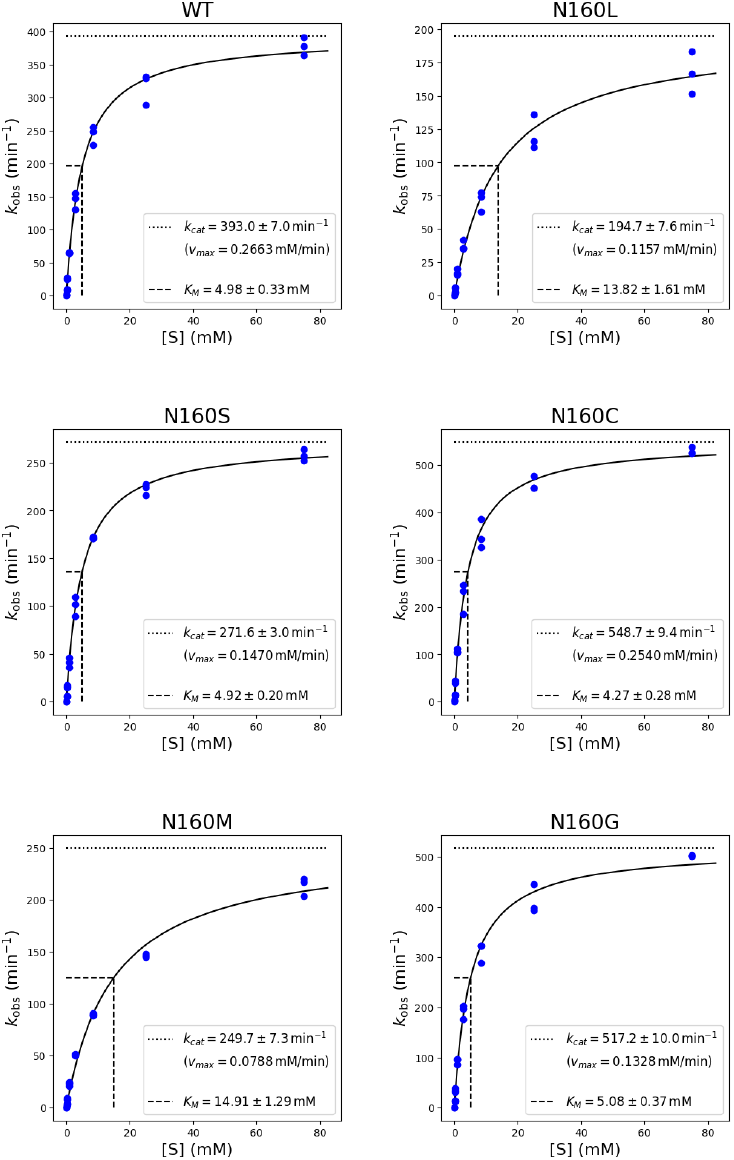
Michaelis-Menten graphs of the wild-type (WT) and five mutants. pPNG concentration [mM] is on the horizontal axis and *k*_obs_ [min^-1^] is on the vertical axis. The triplicate data was fit with nonlinear regression.

### Thermal Stability

Thermal stability was evaluated for the wild-type and all variants using the Protein Thermal Shift assay, performed in triplicate to obtain mean T_M_ values with associated uncertainty (Figure 5). The wild-type enzyme exhibited a T_M_ of 45.9 ± 0.2 °C. Accounting for uncertainty, T_M_ values ranged from 42.4 to 46.7 °C across all samples. Linear regression analysis between T_M_ and TSE produced an R^2^ value of 0.65 with a one-sided p-value of 0.03; the estimated regression line was Y = −0.26X − 242.13 (Figure 5). Simultaneous pairwise confidence interval analysis identified statistically significant differences between N160L and N160G, N160C and N160M, N160C and N160G, and N160G and wild-type (Table 2). At the 95% confidence level, the T_M_ of N160C exceeded that of N160M and N160G by between 1.0 and 5.3 °C and between 1.2 and 5.4 °C, respectively. The T_M_ values for N160L and wild-type were greater than that of N160G by between 1.0 and 3.5 °C and between 1.3 and 4.5 °C, respectively (Table 2).

**Table 2.** Bonferroni-corrected pairwise confidence intervals (α = 0.05) for T_M_ comparisons among wild-type and mutant BglB variants. Welch’s t-test assuming unequal variances was used to calculate intervals. Statistically significant comparisons are shown in bold.

| Comparison | Mean Difference | t-critical | Lower Bound | Upper Bound | Significant |
| --- | --- | --- | --- | --- | --- |
| N160L vs N160S | 0.5 | 7.1 | -0.8 | 1.8 | No |
| N160L vs N160C | 1.5 | 21.9 | -2.5 | 5.4 | No |
| N160L vs N160M | 2.1 | 9.9 | -0.1 | 4.3 | No |
| **N160L vs N160G** | **2.2** | **7.6** | **1.0** | **3.5** | **Yes** |
| N160L vs WT | -0.7 | 7.5 | -1.4 | 0.1 | No |
| N160S vs N160C | -2.0 | 10.2 | -4.1 | 0.0 | No |
| N160S vs N160M | 1.1 | 7.0 | -0.6 | 2.9 | No |
| N160S vs N160G | 1.3 | 6.3 | -0.0 | 2.5 | No |
| N160S vs WT | -1.6 | 10.8 | -3.3 | 0.0 | No |
| **N160C vs N160M** | **3.2** | **8.6** | **1.0** | **5.3** | **Yes** |
| **N160C vs N160G** | **3.3** | **10.3** | **1.2** | **5.4** | **Yes** |
| N160C vs WT | 0.4 | 62.6 | -9.3 | 10.0 | No |
| N160M vs N160G | 0.1 | 7.0 | -1.6 | 1.9 | No |
| N160M vs WT | -2.8 | 13.2 | -5.6 | 0.1 | No |
| **N160G vs WT** | **-2.9** | **10.7** | **-4.5** | **-1.3** | **Yes** |

**Figure 5.**
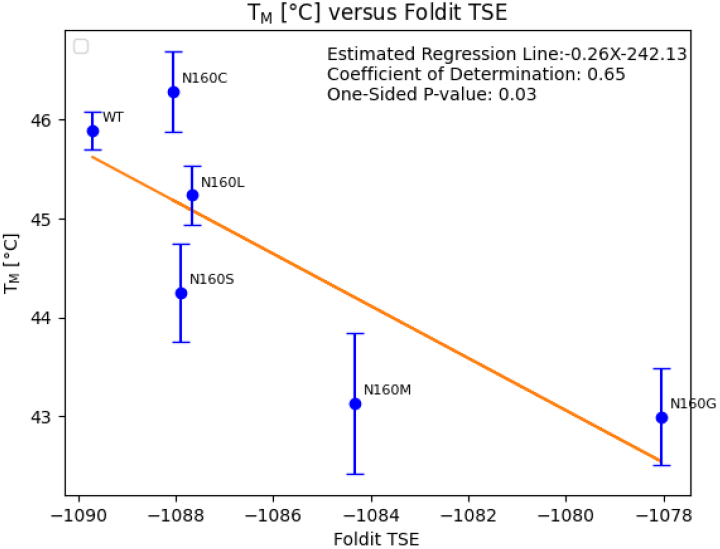
Foldit Standalone TSE scores plotted against T_M_ values shown with error bars [°C] for the wild-type and each mutant. Error is reported as ± 2s (estimated sample standard deviation from t-distribution).

## DISCUSSION

This study examined the catalytic efficiency and thermal stability of five BglB variants, each containing a single point mutation at residue N160. The mutants were selected to represent a range of biophysical properties and to evaluate the contribution of this residue to the enzyme’s kinetic performance and thermal stability.

### Protein Purity and Expression

Despite the loss of the hydrogen-bonding network observed in the wild-type, all mutants were expressed as soluble proteins with concentrations ranging from 1.6 to 3.5 mg/mL.

### Kinetic Activity

The N160L and N160M variants exhibited reduced catalytic performance relative to the wild-type. Both mutants showed approximately 1.5-fold lower *k*_cat_ values and 2.5-fold higher K_M_ values, resulting in more than a 4.7-fold decrease in catalytic efficiency (*k*_cat_/K_M_). This reduction likely arises from the bulky side chains of leucine and methionine, which may intrude into adjacent residue space and restrict local conformational flexibility. Such steric hindrance could impede the conformational changes required for efficient substrate binding and catalysis. Foldit Standalone modeling supports this interpretation, with clashing scores of 5.4 and 15.0 for N160L and N160M, respectively, compared with 0.8 for the wild-type. These elevated scores indicate greater steric interference with neighboring residues.

In contrast, the N160C and N160G variants exhibited improved catalytic performance from the wild-type. Specifically, N160C and N160G showed more than 1.3-fold greater *k*_cat_ values and approximately 0.8-fold lower K_M_ values compared with the wild type. Consequently, their catalytic efficiencies, expressed as *k*_cat_ divided by K_M_, were 1.6-fold and 2.1-fold higher, respectively. The improved catalytic efficiency relative to the wild-type likely reflects the smaller size of cysteine and glycine side chains compared with that of asparagine. Reduced side-chain volume decreases steric constraints, providing additional space for neighboring residues. Based on biophysical principles and our structural model, N160C and N160G appear to impose less side-chain rigidity than N160L and N160M, perhaps permitting greater local mobility that accommodates tighter substrate binding and faster catalysis. Foldit Standalone modeling supports this interpretation, as the side-chain scores quantify each residue’s contribution to predicted folding stability, where lower values are more favorable. N160C and N160G had scores of 0.1 and 0, respectively, compared with 2.4 for asparagine. The Foldit side-chain contribution profiles for N160C, N160G, and the wild-type were consistent with the kinetic data, where lower side-chain contribution scores corresponded to higher catalytic efficiency.

Despite the similar hydrophilic and polar properties of serine and asparagine, the N160S mutant showed reduced kinetic performance compared to wild-type. N160S exhibited 0.6-fold lesser values for both *k*_cat_ and overall catalytic efficiency. We hypothesize this difference is due to the distinct hydrophilic properties of serine versus asparagine. Wherein, the hydroxyl group of serine may alter the water molecule arrangements in the active site required for the substrate reaction (Figure 1b); thus, disrupting the active site geometry and reducing catalytic efficiency. Furthermore, the loss of hydrogen bonding in N160S decreases the structural integrity of the residue site, also affecting the precise coordination required for efficient substrate turnover.

### Thermal Stability

The simultaneous pairwise confidence intervals were constructed to identify statistically significant differences in T_M_ while controlling for multiple comparisons. The analysis showed that the T_M_ of N160C was significantly greater than those of N160M and N160G. In addition, both N160L and the wild-type had significantly greater T_M_ values, in this sample set, than N160G.

Despite previous findings reporting a negligible Pearson correlation (r < −0.001) between T_M_ and TSE ^12^, we found a negative linear relationship between T_M_ and TSE in these data (Figure 5). Building off this preliminary result, we think that certain mutation sites may produce stronger associations between T_M_ and TSE, perhaps making them particularly informative for predictive modeling. It would be interesting to investigate other residue locations that appear to similarly disrupt hydrogen bonding and water arrangements to look for more compelling trends between stability and TSE. Given the independent nature of kinetic and T_M_ parameters ^12^, we wonder if this might be similarly true for a special set of sites wherein model outputs (e.g., TSE) are indeed predictive of catalytic performance.

## CONCLUSION

This study presents a novel contribution to the BglB D2D dataset through the kinetic and thermal characterization of five single-point mutants at residue N160, three of which have not been previously reported. By integrating new experimental measurements with computational parameters from Foldit, this study provides initial evidence of a measurable relationship between total system energy and thermal stability at this residue site. These findings highlight the value of more deeply characterizing select sites to uncover predictive features that influence enzyme performance. Expanding datasets with studies of this kind will improve the accuracy and reliability of predictive models, particularly for enzyme systems where experimental data remain limited. Continued generation of high-quality, functionally relevant data will enhance model development and advance applications in protein therapeutics, biofuels, food science, and bioremediation, bringing us one step closer to the full potential of data-driven enzyme design.

## Acknowledgments

This work was supported by the University of California Davis, the National Science Foundation (award nos. # 2315767, 2118138 and 1827246). The content is solely the responsibility of the authors and does not necessarily represent the official views of the National Science Foundation, or UC Davis. Further, this work references and is contextualized by the Design to Data (D2D) dataset, which has been built through contributions of over 1000 undergraduate students over the past decade.

